# Localized assembly for long reads enables genome-wide analysis of repetitive regions at single-base resolution in human genomes

**DOI:** 10.1101/2022.12.02.518938

**Authors:** Ko Ikemoto, Hinano Fujimoto, Akihiro Fujimoto

## Abstract

**Background:** Long-read sequencing technologies have the potential to overcome the limitations of short reads and provide a comprehensive picture of the human genome. However, it remains hard to characterize repetitive sequences by reconstructing genomic structures at high resolution solely from long reads. Here, we developed a localized assembly method (LoMA) that constructs highly accurate consensus sequences (CSs) from long reads.

**Methods:** We first developed LoMA, by combining minimap2, MAFFT, and our algorithm, which classifies diploid haplotypes based on structural variants and constructs CSs. Using this tool, we analyzed two human samples (NA18943 and NA19240) sequenced with the Oxford Nanopore sequencer. We defined target regions in each genome based on mapping patterns and then constructed a high-quality catalog of the human insertion solely from the long-read data.

**Results:** The assessment of LoMA showed high accuracy of CSs (error rate < 0.3%) compared with raw data (error rate > 8%) and superiority to the previous study. The genome-wide analysis of NA18943 and NA19240 identified 5,516 and 6,542 insertions (≥ 100 bp) respectively. Most insertions (∼80%) were derived from the tandem repeat and transposable elements. We also detected processed pseudogenes, insertions in transposable elements, and long insertions (> 10 kbp). Further, our analysis suggested that short tandem duplications were association with gene expression and transposons.

**Conclusions:** Our analysis showed that LoMA constructs high-quality sequences from long reads with substantial errors. This study revealed the true structures of insertions with high accuracy and inferred mechanisms for the insertions. Our approach contributes to the future human genome studies. LoMA is available at our GitHub page: https://github.com/kolikem/loma.

## Background

Long-read sequencing technologies, such as Oxford Nanopore Technologies (ONT) and Pacific Biosciences (PacBio), have the potential to overcome the limitations of next-generation sequencing technologies and provide a comprehensive picture of the human genome [1]. Current technological shortcomings, such as high sequencing error rates, are steadily being overcome, and many recent studies using long reads have given new findings in previously inaccessible regions [2][3][4]. Further, using long reads, they identified pathogenic structural variants (SVs) such as large deletions in *EYS* in retinitis pigmentosa [5] and the expansion of a triplet repeat in *NOTCH2NLC* in neuronal intranuclear inclusion disease [2]. Moreover, there is a growing demand for clinical applications of long reads studies to solve common and Mendelian disorders [1]. Studies using long reads may thus contribute to new therapeutics and more accurate diagnosis of human disease.

However, many problematic regions are still harbored in the human genome. Widely distributed repeats, such as tandem and interspersed repeats, can cause issues in the data processing [6]. Expansions of tandem repeats (TRs) cause various disorders [2][7][8], and transposable elements (TEs) including *Alu* and LINE elements have been associated with a range of disorders [9][10]. Thus, the analysis of these problematic regions is a very important for the study of human genetics.

Recently, the first complete sequence of a human genome was finished by the Telomere-to-Telomere (T2T) Consortium (T2T-CHM13) [11], which added 238 Mbp of non-syntenic sequences to the GRCh38 assembly. This study also showed that most additional bases were derived from repetitive sequences, such as centromeric satellites and segmental duplications, suggesting that previous research based on *de novo* assembly may have dismissed a large portion of repetitive sequences. Although the T2T-CHM13 assembly is one of the greatest achievements of long reads, it was attained by sequencing the cell line from a complete hydatidiform mole (uniformly homozygous) and by combining data from multiple platforms including HiFi, ONT, Illumina, and other state-of-the-art techniques. Such large-scale, comprehensive approaches cannot be applied to practical clinical studies in most cases due to costs and human resources.

Current SV calling is essentially based on two types of methodologies: a *de novo* assembly-based approach and a mapping-based approach [12]. One of the advantages of the former lies in the detection of large variants [12][13]. However, this method is hampered by haplotype representations and assembly errors caused by repetitive sequences [14]. On the other hand, a mapping-based approach with long reads is advantageous when coverage of the available data is low or the samples contain low-frequency SVs [12]. The variant discovery of this approach usually begins with the mapping of reads to a reference genome and detects SVs based on the mapping patterns. However, misalignments due to sequencing errors sometimes cause false positives and negatives, and tangled, nested SVs prevent accurate SV calling. Considering these advantages and disadvantages of the current approaches, we aimed to establish a hybrid approach to analyze SVs at single-base resolution.

Therefore, here we developed a localized assembly tool, LoMA (Localized Merging and Assembly), which generates accurate consensus sequences (CSs) based on the mapping results of long reads. LoMA captures haplotype structures based on SVs and produces haplotype-resolved CSs, which help identify heterozygous variants. To our best knowledge, only one tool, lamassemble, has a similar function [15]; however, this software cannot classify heterozygous regions, which can hinder the resolution of human diploid genomes. In response, we implemented a haplotype classification of target regions which improved the accuracy of SV detection.

We applied LoMA to genome-wide SV detection in two samples using the high-coverage whole-genome sequencing (WGS) data from ONT. Generally, true genomic structures of insertions are difficult to resolve. Thus, we aimed to reveal true inserted sequences at single-base resolution. Our analysis showed that LoMA constructs high-quality sequences from single-platform data with substantial errors and revealed the true structures of tandem and interspersed repeats in the human genome.

## Methods

### Development of LoMA

Raw reads yielded by ONT are accompanied by substantial sequencing errors. LoMA is a tool that assembles long reads mapped to a certain region and generates a highly accurate CS (Fig. 1A, B). LoMA detects heterozygous SVs in a target region and outputs haplotype-resolved sequences. In the first step, LoMA constructs a CS spanning a target region. This process is initiated by finding overlaps of raw reads using pairwise all-to-all alignment of minimap2 (`-x ava-ont`) [16], followed by a layout of overlapped reads (Fig. 1A). A read layout is then divided into blocks, and raw reads in every block undergo a multiple alignment using MAFFT [17]. Based on the multiple alignment, each consensus nucleotide is determined at every position, and partial CSs are generated for each block (Fig. 1A). By concatenating the series of short blocks, one complete CS that covers the whole target region is obtained (Fig. 1A).

**Fig. 1:**
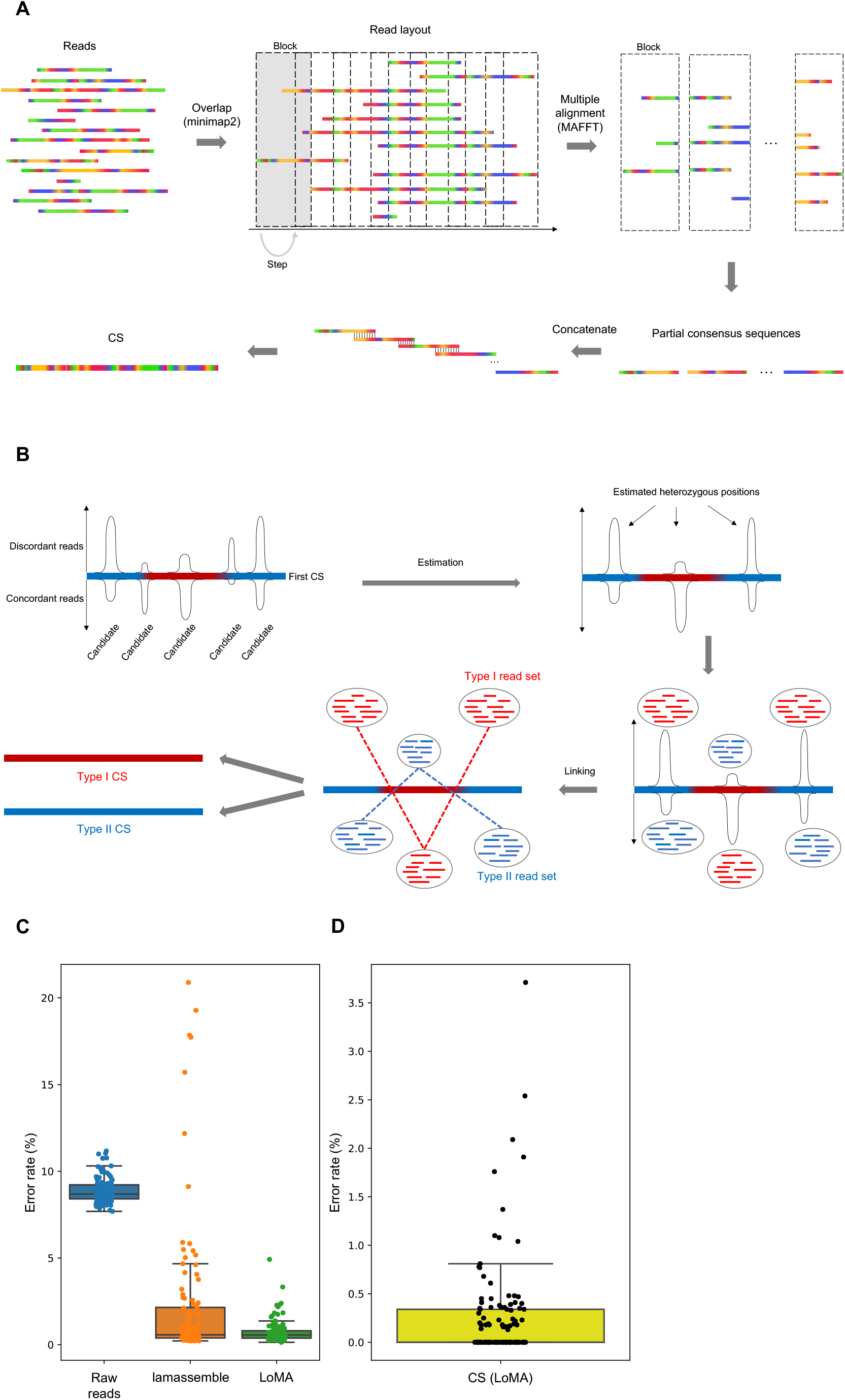
Schema and accuracy assessment of LoMA. **A**. The main process to compose a consensus sequence (CS) is shown. Reads are laid out based on an all-to-all alignment using minimap2 [16]. The layout is then subdivided into short blocks for the successive procedure to determine partial consensus sequences based on a multiple alignment using MAFFT for each block [17]. A CS is obtained by concatenating these short blocks. **B**. The read classification process is shown. Reads constituting a CS are aligned to the CS itself using minimap2 [16]. LoMA predicts heterozygous loci in the region based on the extent of deviation from the binomial distribution, and the reads derived from each estimated haplotype are gathered (Type I and II read sets). Both sets return to the main process (A) after this step. **C**. A comparison with raw reads (blue) and sequences by lamassemble (orange). Consensus sequences (CSs) by LoMA (green) showed a sharp drop in the error rate compared with raw reads and lamassemble. The standard deviations are 0.67, 4.09 and 0.72 for LoMA, lamassemble and raw reads respectively, which shows the dispersion of LoMA is smaller than of lamassemble, and the instability of sequences in raw data is reduced by LoMA. **D**. A comparison of CSs constructed by LoMA with sequences by Sanger sequencing. The sequences from 121 insertions of NA18943 showed high accuracy (99.7% at least). Individually, 66 out of 121 (54.5%) regions constructed by LoMA were error-free compared with poly(A) compressed sequences.

The generated CS can be a mosaic sequence mixing paternal and maternal haplotypes, resulting in the dismissal of one allele of the heterozygous variants. To overcome this problem, we adopted the read-separation step (Fig. 1B). In this step, all input reads are aligned to the first CS recurrently by using minimap2 [16]. Then, bins (700 bp) with mismatch clusters are detected based on two conditions: the number of discordant reads to the first CS > 7; and the number of discordant reads is within the mean ± 3σ of a binomial distribution with *p* = 0.5. A pair of haplotypes that have two heterozygous SVs is separated using reads that span both SVs (Fig. 1B). By tracking back all heterozygous bins in a target region, all reads are coherently classified into two individual read sets (Type I and II) in most cases (Fig. 1B). CSs are generated from both groups of reads. If a target region contains no SVs, two read sets are not produced and the region is regarded as homozygous.

### DNA sequencing and data processing

Two samples, NA18943 and NA19240, from the International HapMap Project was analyzed [18]. NA19240 is a Yoruban male and was sequenced with PromethION (ONT) by De Coster et al. [19]. The WGS data of this sample were downloaded in FASTQ format from the European Nucleotide Archive (accession number: PRJEB26791) [20]. NA18943 is a Japanese male and the first Japanese sample sequenced by short reads [21]. Its genomic DNA was extracted from a B cell line. In total, 26 runs were performed using MinION (ONT).

Using the WGS data of NA18943, basecalling was performed using Guppy (ver.4.4.1) [22]:

`--flowcell FLO-MIN106 --kit SQK-LSK109 --device auto`

The sequencing data of both NA18943 and NA19240 were then aligned to the GRCh38 assembly using minimap2 (ver.2.0-r290-dirty) [16]:

`-a -g2000 -A1 -B2 -O2,32 -E1,0 -z200`

### Accuracy assessment of LoMA

To estimate the accuracy of LoMA, we assessed CSs by comparing them with GRCh38. We randomly selected 108 positions from the human genome, excluding the centromeres and gaps (Additional file 1: Table S1). Using the data of NA18943, we collected all reads mapped within 20 kbp of each position and constructed CSs using LoMA. We then aligned the generated CSs to GRCh38 using minimap2 [16] and calculated the error rate from the CIGAR strings in SAM files. Similarly, we aligned all reads collected from the 108 regions to GRCh38 and calculated the error rate for each read. Also, we assembled matched regions based on the same sets of reads using lamassemble for benchmarking [15]:

`-P 8 -a -v -p 2e-3 -m 2*(number of reads) -z 1000 promethion.mat`

The error rate of lamassemble was calculated in the same manner.

### Detecting unclear regions in genomes

To detect SVs from the two samples, we applied LoMA to the WGS data. We first searched for target regions (unclear regions) by scanning all chromosomes from telomere to telomere. We split each autosome and sex chromosome binned per 500 bp, step size 250 bp, and defined an “unclear” region as follows: (i) average coverage between 10 and 200, (ii) total number of reads containing indels (100 bp) or hard-or soft-clipped sequences (500 bp) > 10, and (iii) the proportion of reads containing indels (100 bp) or hard-or soft-clipped sequences (500 bp) > 0.2 (Fig. 2A). Then, the bins within 10 kbp of each other were merged together. We defined the merged bins as unclear regions in this study. After determining the unclear regions for each genome, we collected all reads that mapped within 10 kbp from the boundary of each region using SAMtools [23].

**Fig. 2:**
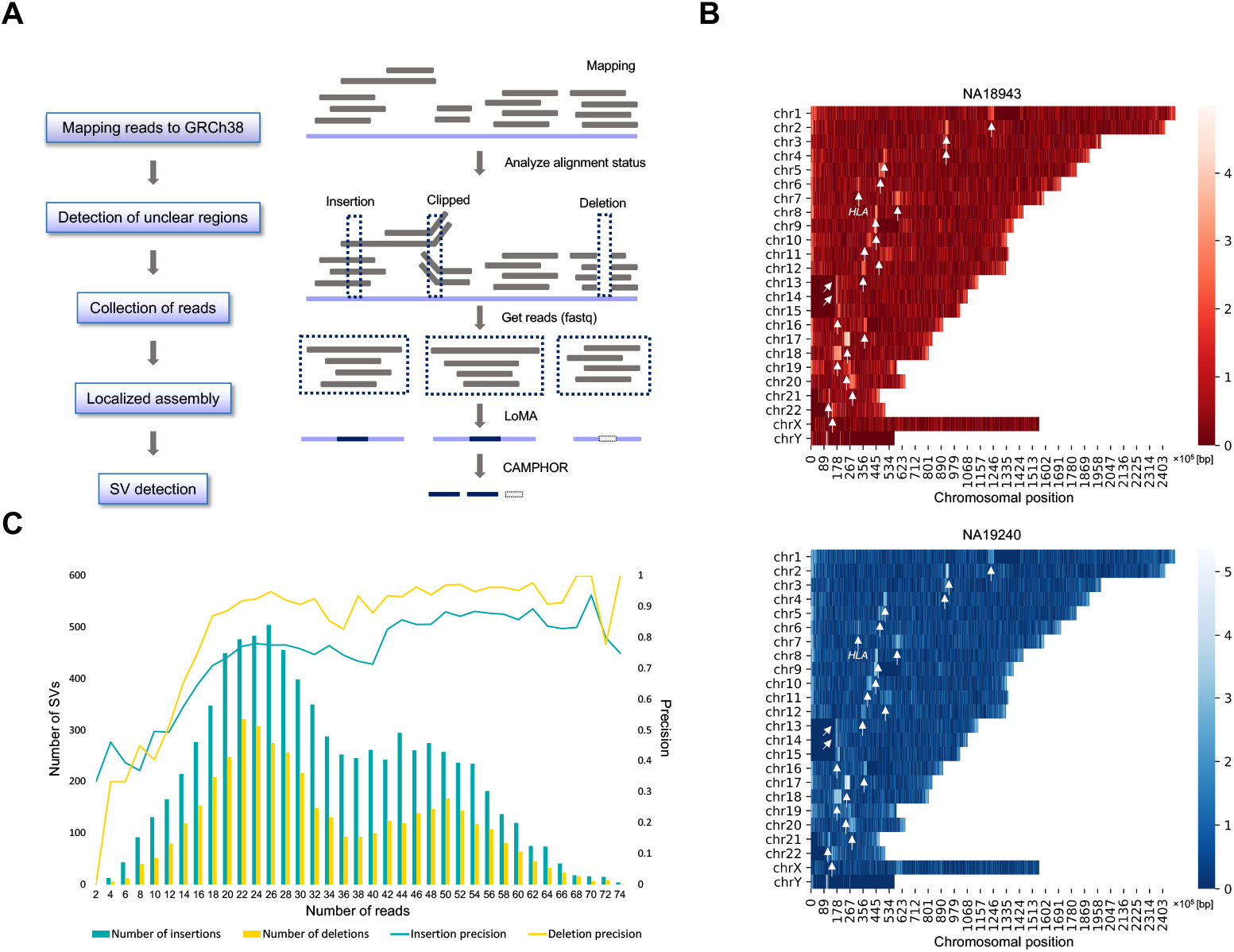
Whole-genome LoMA analysis. **A**. The workflow of the whole-genome analysis. Unclear regions were first defined based on the alignment status (indels and clips) of ONT reads. Reads mapped to the regions were separately collected. For each region, LoMA attempted localized assembly to obtain accurate sequences. CAMPHOR collectively detected variants (≥ 20 bp) from the CSs for NA18943 and NA19240. **B**. The relative density of unclear regions of NA18943 and NA19240 are shown in a red-and blue-colored heatmaps, respectively. The light-colored regions are dense with unclear regions. The white arrows indicate the autosomal centromeres except chromosome 6. The arrows on chromosome 6 represent the *HLA* region. **C**. The precision of indels to the standard SV set was assessed for NA19240. The left vertical axis (bar graph) shows the number of indels found at each bin (the number of constitutive reads). The right (line graph) shows the precision of indels at each bin. Both graphs are binned per 2 reads.

### Localized assembly and SV detection

To obtain CSs from the defined unclear regions above, we analyzed FASTQ files using LoMA and constructed CSs with the following parameters.

`block=3000 step=1000 hashicut=10 sigma=3`

We then called indels (≥ 100 bp) found in the CSs using CAMPHOR [3].

### Experimental validation and benchmark of SV detection

To evaluate the accuracy of inserted sequences obtained from CSs, we performed Sanger sequencing. We randomly selected 121 homozygous insertions from NA18943. PCR primers were designed in flanking regions, and the PCR-direct sequence was conducted. We compared the sequences by Sanger sequencing with the matched sequences of CSs generated by LoMA. Since the length of homopolymers was difficult to determine by Sanger sequencing, the stretches of poly(A) were removed from the comparison.

An accurate SV callset of NA19240 was released (the standard SV set) in a previous study leveraging multi-platform data [24]. We compared the indels of our SV calling result (the LoMA SV set) with those of the standard SV set and assessed the precision. We considered variants from the LoMA SV set as concordant (true positive) with the standard SV set if both were the same type and the distance between two breakpoints of the SVs was < 500 bp.

### Decomposing insertions

Insertion events are known to be caused by various mechanisms and have various consequences [25]. To characterize and investigate the origins of the detected insertions, we decomposed them into TRs, TEs, tandem duplications (TDs), satellite sequences, dispersed duplications, processed pseudogenes, alternative sequences, “deletions” in GRCh38, and nuclear mitochondrial DNA sequences (NUMTs).

We first applied Tandem Repeats Finder (TRF) [26] to all inserted sequences and defined TRs as having (i) element lengths < 50 bp and (ii) covering more than 50% of an inserted sequence. After filtering TRs, we identified TEs using RepeatMasker [27] if (i) an inserted sequence covered a TE > 50%, (ii) the inserted sequence was covered by the TE > 50% (reciprocal overlap), and (iii) the total substitutions and indels < 50% (matching condition).

Previous studies have reported TDs that are still understudied but widespread [25][28]. After detecting TRs and TEs, we manually reviewed the remaining insertions and found that they contained TDs derived from nonrepetitive regions in the reference. We considered these insertions as TDs. To identify this class of insertions, we aligned all insertions except TRs to GRCh38 using BLAT [29]. We then collected insertions mapped to original breakpoints within 5 bp with > 90% in BLAT identity and defined them as TDs. In this process, missing TRs with long repeat elements were also found. Therefore, they were added to the TR callset if (i) an inserted sequence aligned within 500 bp from the insertion breakpoint and (ii) the ratio of the total number of matching bases to the insertion length > 0.5.

To understand the remaining insertions, we manually checked their features by aligning them to the reference using BLAT [29]. We identified insertions that were aligned from end to end to different chromosomal regions with high identity (> 90%). We defined these insertions as dispersed duplications. Next, we detected insertions aligned to a series of exons and untranslated regions (UTRs) from coding genes with high identity (> 90%) and classified them as processed pseudogenes. We also found other insertions aligned to the alternative sequences (e.g., “alt” or “fix” sequences) on BLAT with high identity (> 90%) and classified them as alternative sequences. Some of the insertions left at this point were thought to have arisen by deletion events in GRCh38 [3], because they were securely aligned to the chimpanzee reference genome (panTro6) even though they became insertions in GRCh38. We aligned the remaining insertions to the panTro6 assembly and categorized the insertions that lifted over panTro6 with high identity (> 90%) within 100 bp from the inserted position on GRCh38 as “deletions” in GRCh38. After this, the remaining insertions were manually reviewed, and features of the genomic regions (segmental duplications or self-chain) were examined.

### Features of tandem repeats

TRs are well known to have a high mutation rate [30]. We investigated the expansion rates of TRs by comparing them with the reference (GRCh38) as follows:

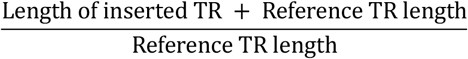

To investigate the composition of expanded elements, we next analyzed repeat element lengths and repeat sequence elements for short tandem repeats (STRs, 2-6 bp). We indiscriminately treated a group of STRs that were the same in lexicographical order without discriminating strands (see Additional file 2). We next analyzed TR expansions in coding genes using the definition of RefSeq gene database [31]. We also analyzed expansions in TEs and surrounding regions. We referred to the repeat annotated GRCh38 sequence by RepeatMasker [27] and counted the number of TR expansions detected inside, upstream (< 100 bp) and downstream (< 100 bp) of TEs. Binomial tests were performed to examine the bias in the frequency of TR expansions upstream and downstream of SINEs.

### Features of target site duplications in Alu element

Several families of TEs, including L1, *Alu* and SVA elements, remain active in the human genome [32]. In the integration process of an *Alu* element, a duplicated sequence is copied and inserted at the flanking site, which is called a target site duplication (TSD) [33]. This is the hallmark of retrotransposition. By surveying *Alu* insertions accompanied by TSDs, we analyzed their characteristics. We made a non-redundant *Alu* insertion set by merging the results using the NA18943 and NA19240 data (merging condition: pairwise distance < 500 bp). We then used the MEME Suite to find motifs at the first and second nicking sites [34]. We also analyzed the length distribution of the TSDs of *Alu* elements.

### Features of tandem duplications

We next explored a class of underrepresented insertions, TD. TD is the duplication of a single copy sequence. However, the mechanism of its generation remains unclear. We analyzed gene expression levels between genes with and without TDs using GTEx data [35]. To examine differences in expression between genes with and without TD, we performed a Wilcoxon rank-sum test. We also performed binomial tests to analyze the association of TDs with TEs based on the repeat annotation of GRCh38 [27] and with coding genes based on the RefSeq gene data [31].

## Results

### Sequencing data

We sequenced the genomic DNA of NA18943 from a B cell line using a single platform, MinION (ONT). After combining yields from 26 flow cells, the sequencing data totaled 231.6 Gbp (77× coverage) (Additional file 1: Table S2). For NA19240, the amount of sequencing data was 255.8 Gbp (79× median coverage) [19].

### Reduction in error rate and accuracy validation of LoMA

To evaluate the accuracy of LoMA, we performed extensive validations. First, we randomly selected 108 regions and constructed CSs (Additional file 1: Table S1). Thirteen of the 108 regions were classified as heterozygous, and two CSs were generated for each heterozygous region. In total, 121 CSs were obtained. We aligned them to GRCh38 and assessed the error rate. The total alignment length was approximately 7.3 Mbp (Additional file 1: Table S3). The estimated error rate was 8.7% (SD = 0.72) for raw reads and 0.76% (SD = 0.67) for CSs by LoMA (Fig. 1C). Similarly, we attempted to construct CSs using lamassemble for a comparison, but it failed to assemble 3 of the 108 regions (Additional file 1: Table S1). A comparison of the successful 105 sequences with GRCh38 estimated the mean error rate of lamassemble to be 2.3% (SD = 4.1) (Fig. 1C).

We next performed an experimental validation for the CSs generated by LoMA. We found 14 SV candidates from 13 heterozygous regions (Additional file 1: Table S1, 4). The primer design and PCR amplification were successful for eight heterozygous candidates, and the product sizes of all candidates were concordant with the expectation of LoMA (Additional file 1: Table S4, Additional file 2: Fig. S1).

Further, we performed Sanger sequencing for 121 homozygous insertions randomly selected from NA18943 (Additional file 1: Table S5). The total compressed length amounted to 66,126 bp, and 65,932 bp were matched to CSs by LoMA (sequence identity 99.71%) (Fig. 1D, Additional file 1: Table S5). Also, 66 CSs from the 121 insertions (54%) were perfectly matched (100%) to the corresponding Sanger sequences (Additional file 1: Table S5). These results suggest that LoMA has good accuracy for the analysis of SVs at single-base resolution.

### Genome-wide analysis of SV polymorphism using LoMA

To detect SV polymorphisms using LoMA, we identified unclear regions based on the number of reads with clipped sequences or indels (Fig. 2A and “Methods”). In NA18943, we detected 27,562 bins as candidates, and they were combined into 16,544 unclear regions. In NA19240, 56,158 candidate bins were detected and combined into 18,928 unclear regions. The larger number of unclear regions in NA19240 reflects the high genetic diversity of African populations [36]. Both NA18943 and NA19240 showed a similar pattern of the landscape of unclear regions: they were localized in the pericentromeric and *HLA* region in both samples (Fig. 2B).

We next constructed the sequences from the unclear regions. After the LoMA analysis, we obtained CSs of 13,822 and 15,220 regions for NA18943 and NA19240, respectively, including 3,065 and 7,542 heterozygous regions (see Additional file 2).

### Comparison of LoMA with a “standard” SV set and threshold for SV detection

The benchmark with a standard SV set of NA19240 indicated that the precision decreased under 20× coverage, although it became stable after more than 20 reads (Fig. 2C). The mean precision of insertions and deletions with 20 or more reads was 0.82 and 0.93, respectively. For less than 20 reads, it was to 0.52 for each. However, the number of SVs with less than 20 reads in the entire SV set was 21.9% for insertions and deletions, respectively; thus, excluding these SVs did not strongly affect the conclusions of this study. Accordingly, we focused on SVs in regions with 20 or more reads to define a more conservative callset of indels. After this filtration, we finally obtained 5,516 insertions and 2,687 deletions in NA18943, and 6,542 insertions and 3,475 deletions in NA19240 (the LoMA SV set).

### Overview of insertions and deletions

The length distributions of the indels were consistent with a previous study (Fig. 3A) [3], suggesting that the genome-wide analysis using LoMA provided a reliable result. Sequence analysis at single-base resolution is effective at obtaining a comprehensive catalog of insertions to infer the biological mechanisms of the insertions. To understand the characteristics of the detected insertions, we first classified them (Fig. 3B, C and “Methods”). The numbers of TRs were 2,841 (51.5%) and 2,922 (44.7%) in NA18943 and NA19240, respectively, TEs totaled 1,819 (33.0%) and 2,506 (38.3%), and TDs amounted to 71 (1.3%) and 88 (1.3%). Approximately 1.5% (NA18943) and 1.2% (NA19240) of the insertions were attributed to variants in satellite sequences.

**Fig. 3:**
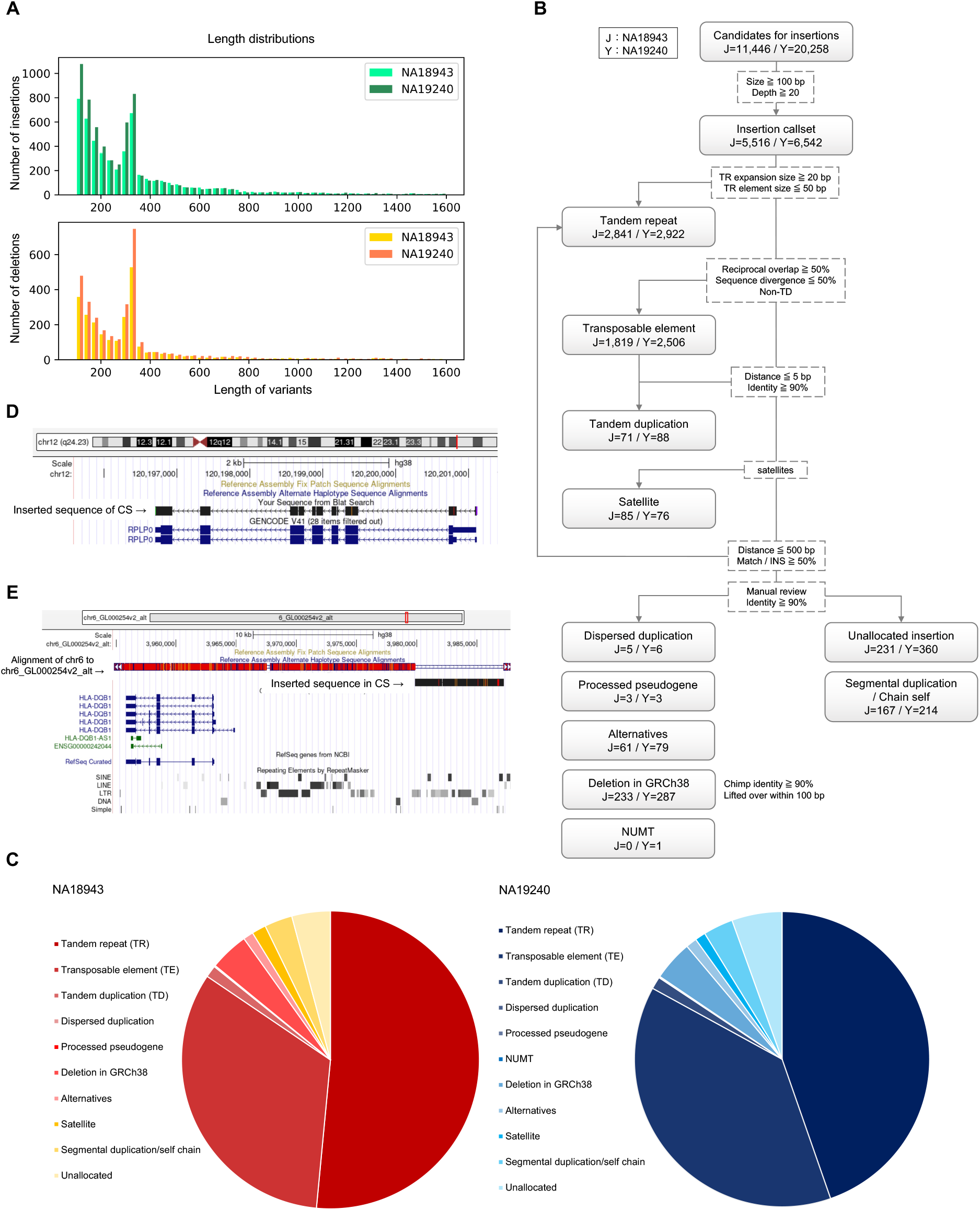
Overview of the variant detection. **A**. The length distributions of insertions (top) and deletions (bottom). The maximum length was capped at 1,600 bp. **B**. The flowchart shows the breakdown of the insertions. Dotted-line boxes represent the conditions of classification, and solid-line boxes are each class of insertions. **C**. The pie charts show the proportion of each class of insertions of NA18943 (left) and NA19240 (right). Insertions were decomposed into the classes: TRs (tandem repeats), TEs (transposable elements), TDs (tandem duplications), dispersed duplications, processed pseudogenes, NUMT, deletions in GRCh38, alternative sequences, satellites, and others. **D**. A processed pseudogene (*RPLP0*) detected in NA19240 is shown (in black). The picture was extracted from Genome Browser. The BLAT identity was 99.9% compared with the reference gene, and the alignment pattern of 5’ UTR showed a closer resemblance with the splicing variant shown at the bottom in the picture. **E**. An insertion detected in NA19240 matching the alternative sequence is shown (in black). The picture was extracted from Genome Browser. The breakpoint was 15 kbp upstream of *HLA-DQB1*. LoMA constructed part of the alternative sequence accurately (99.8% in BLAT identity).

We manually reviewed the remaining insertions and identified other types of insertions (Fig. 3B, C): three processed pseudogenes from each of NA18943 and NA19240, five and six dispersed duplications, respectively, and 61 (1.1%) and 79 (1.2%) alternative sequences, respectively. For example, a processed pseudogene found in NA19240, sized 1,115 bp (the breakpoint was at chr11:60,274,156), was precisely mapped to *RPLP0* on chromosome 12 with 99.9% in BLAT identity (Fig. 3D). This processed pseudogene seemed to represent a specific splicing variant. Notably, we found 233 (4.2%) and 287 (4.4%) variants in NA18943 and NA19240 mapped to panTro6, although they did not exist in GRCh38 (“deletions” in GRCh38), suggesting that they should have been caused by deletion events in the genome constituting the GRCh38 assembly [3]. We also found one NUMT insertion, sized 532 bp in NA19240.

After our manual review, 398 (7.2%) and 574 (8.8%) insertions remained to be assigned in NA18943 and NA19240, respectively (Fig. 3B, C). Of those insertions, 167 (3.0%) and 214 (3.3%) occurred in segmental duplications and self-chains in NA18943 and NA19240, respectively. The numbers of unallocated insertions were 231 (4.2%) and 360 (5.5%) (Fig. 3B).

The insertions mapped to alternative sequences tended to be large (median length = 4,049 bp). In NA19240, 12 insertions (15%) mapped to the alternative sequences were not found in the standard SV set (Additional file 1: Table S6). For instance, an insertion sized 7,332 bp, derived from an alternative sequence (chr6_GL000254v2_alt), was found in the LoMA SV set (the breakpoint was chr6:32,687,972). This breakpoint was located 15 kbp upstream from *HLA-DQB1* in the *HLA* region and showed a high sequence identity (99.8%) to chr6_GL000254v2_alt (Fig. 3E). However, unlike an insertion, a translocation was identified in the standard SV set near this region, suggesting an error of the previous SV calling [24]. In NA18943, we found the longest insertion, sized 14,330 bp, at chr12:127,153,629 derived from an alternative sequence (chr12_KZ559112v1_alt). This insertion also showed a high sequence similarity (99.8% in identity) to the alternative sequence (Additional file 2: Fig. S2).

### Tandem repeats (TRs)

Most of the TR insertions were found in TRs in GRCh38 in NA18943 and NA19240. An analysis of expansion rates showed that most TRs had low expansion rates (Fig. 4A), but high expansion rates were observed mainly in short TRs (< 100 bp) (Fig. 4A). An analysis of repeat elements showed that expansions of triplet repeats were rarer than 4-and 5-bp unit repeats in NA18943 and NA19240 (Fig. 4B). Expansions of (AT)*n* were major in 2-bp unit TRs. In 3-, 4-, and 5-bp unit repeats, (AGG)*n*, (AGGG)*n* and (AGGGG)*n* were dominant, respectively (Fig. 4C). In both samples, 42% (NA18943) and 39% (NA19240) of TRs were found in genic regions, which is consistent with the size of the genic region in the human genome (Additional file 2: Fig. S3). Only 7 and 12 genes contained exonic TR expansions in NA18943 and NA19240, respectively (Additional file 1: Table S7). However, they were associated with various human traits and disease, such as the telomere length (Additional file 1: Table S7).

**Fig. 4:**
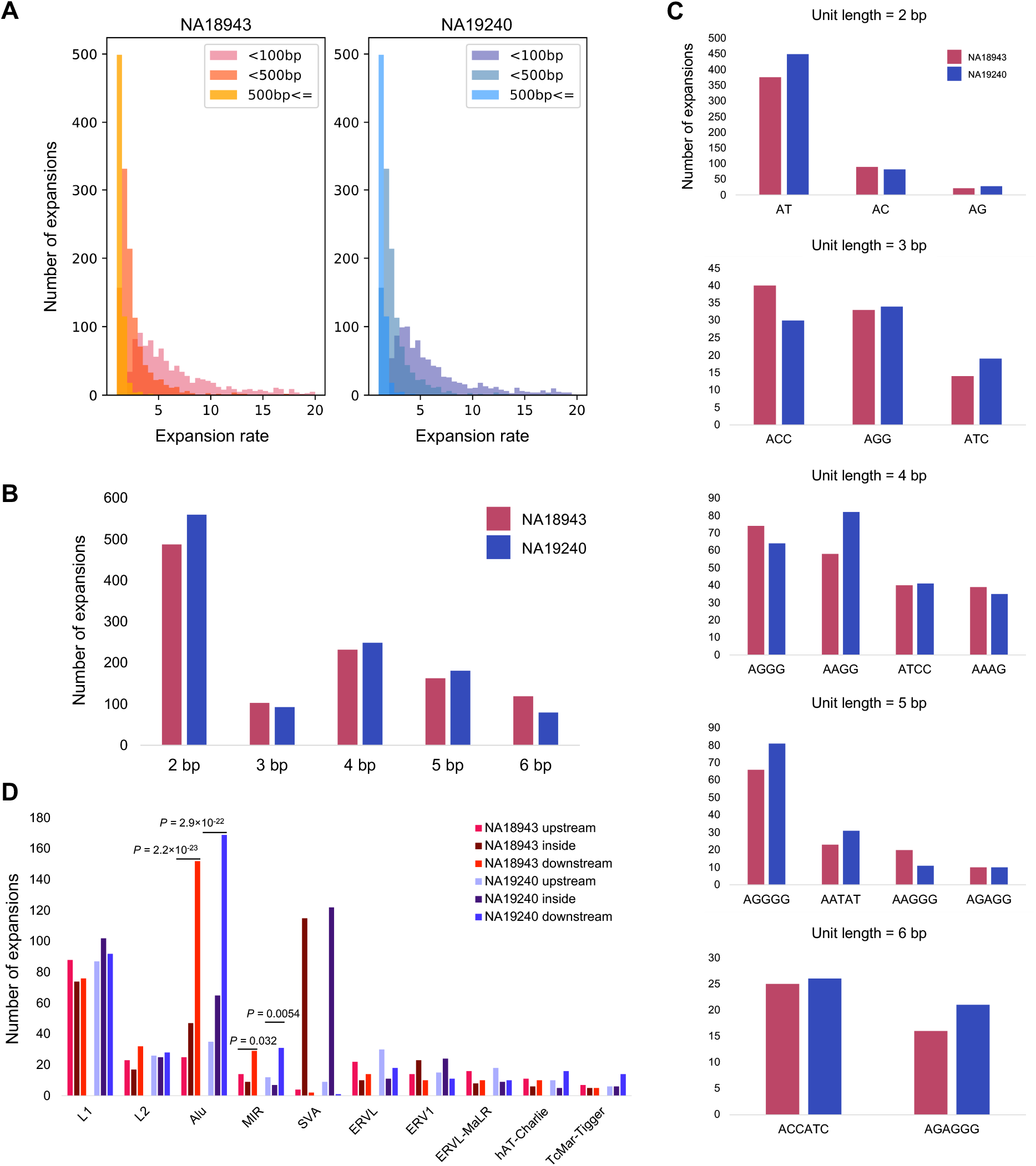
The variation of TR expansions. **A**. The expansion rate is shown up to 20. The left panel, NA18943; the right panel, NA19240. Both panels are separated by reference TR length (< 100 bp, < 500 bp, ≥ 500 bp). **B**. The numbers of variants of STRs (2-6 bp) for NA18943 and NA19240. **C**. The patterns of STR expansions are shown individually by unit length for NA18943 and NA19240. Elements in both samples with a frequency of at least 10 are displayed. **D**. Sites of the occurrence of TR expansions surrounding TEs. NA18943 is red and NA19240 blue; each class is separated into the upper stream (< 100 bp), lower stream (< 100 bp) and inside from the left. In SINE (*Alu* and *MIR*), TR expansions often occurred downstream of TE elements. SVAs contained relatively many expansions inside considering the fewer number than in the LINE family.

We also analyzed the association between TRs and TEs. We found that TR expansions around TEs were more likely to occur downstream of SINEs (*Alu* and *MIR*), than upstream (binomial test: *P* = 2.2 × 10^−23^ (*Alu*) and *P* = 0.032 (*MIR*) in NA18943; and *P* = 2.9 × 10^−22^ (*Alu*) and *P* = 0.0054 (*MIR*) in NA19240) (Fig. 4D). However, we did not see the same tendency in LINEs (L1 and L2) (Fig. 4D). In SINE-VNTR-*Alu* (SVA) elements, many TR expansions were observed inside compared with other families, such as LINEs, possibly because of the unique structure of SVA elements containing a VNTR region (Fig. 4D).

### Target site duplications (TSDs) in Alu elements

In NA18943 and NA19240, 1,579 non-redundant *Alu* insertions were detected. Our motif search of flanking sequences detected a strong motif at the first nicking site, TTAAAAA (*E*-value = 2.5 × 10^−315^) (Fig. 5A), a result consistent with other studies [32][33][37]. However, no clear motif was identified around the second nicking site. We next analyzed the length distribution of TSDs flanking *Alu* elements, and single-peak distributions (median length = 15 bp) were obtained (Fig. 5B). The overall shape of the distributions was slightly skewed left, showing TSDs longer than 15 bp were fewer than TSDs shorter than 15 bp.

**Fig. 5:**
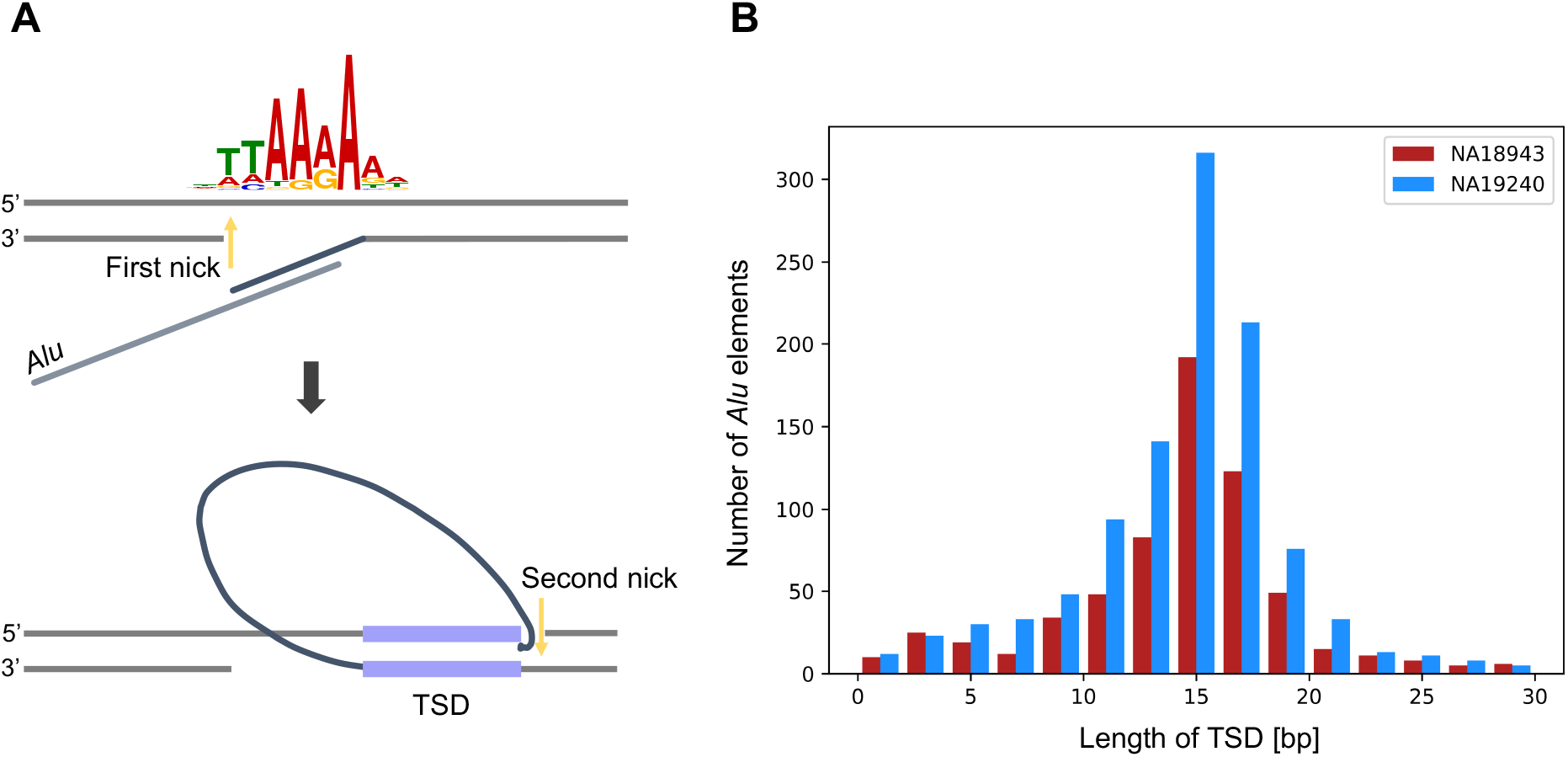
Analysis of TSDs among *Alu* insertions. **A**. The motif search using 1,579 *Alu* elements by MEME [30] confirmed the motif previously studied at the first nicking site. **B**. The length distribution of TSDs among *Alu* insertions, capped at 30 bp, binned per 2 bp. The distribution had a peak around 15 bp and was symmetrical in both samples.

### Tandem duplications (TDs)

Lastly, we detected and analyzed TDs (*n* = 71 in NA18943 and *n* = 88 in NA19240). The length distribution showed that TDs were at most ∼600 bp in length, with shorter TDs dominant (Fig. 6A). A manual review using UCSC Genome Browser [38] showed that 50 (NA18943) and 54 (NA19240) TDs were observed in TEs and significantly enriched in TEs in NA18943 and NA19240, respectively (binomial test: *P* = 1.3 × 10^−5^ in NA18943 and *P* = 4.0 × 10^−3^ in NA19240) (Fig. 6B). We next analyzed the association of TDs and genes. TDs were observed in 30 (NA18943) and 34 (NA19240) genes, and together, 54 non-redundant TDs in genic regions were identified (Additional file 1: Table S8). The genes with TDs showed higher expression levels than other genes (Wilcoxon test: *P* = 4.4 × 10^−12^) (Fig. 6C), although TDs were not enriched in gene regions (Additional file 2: Fig. S4).

**Fig. 6:**
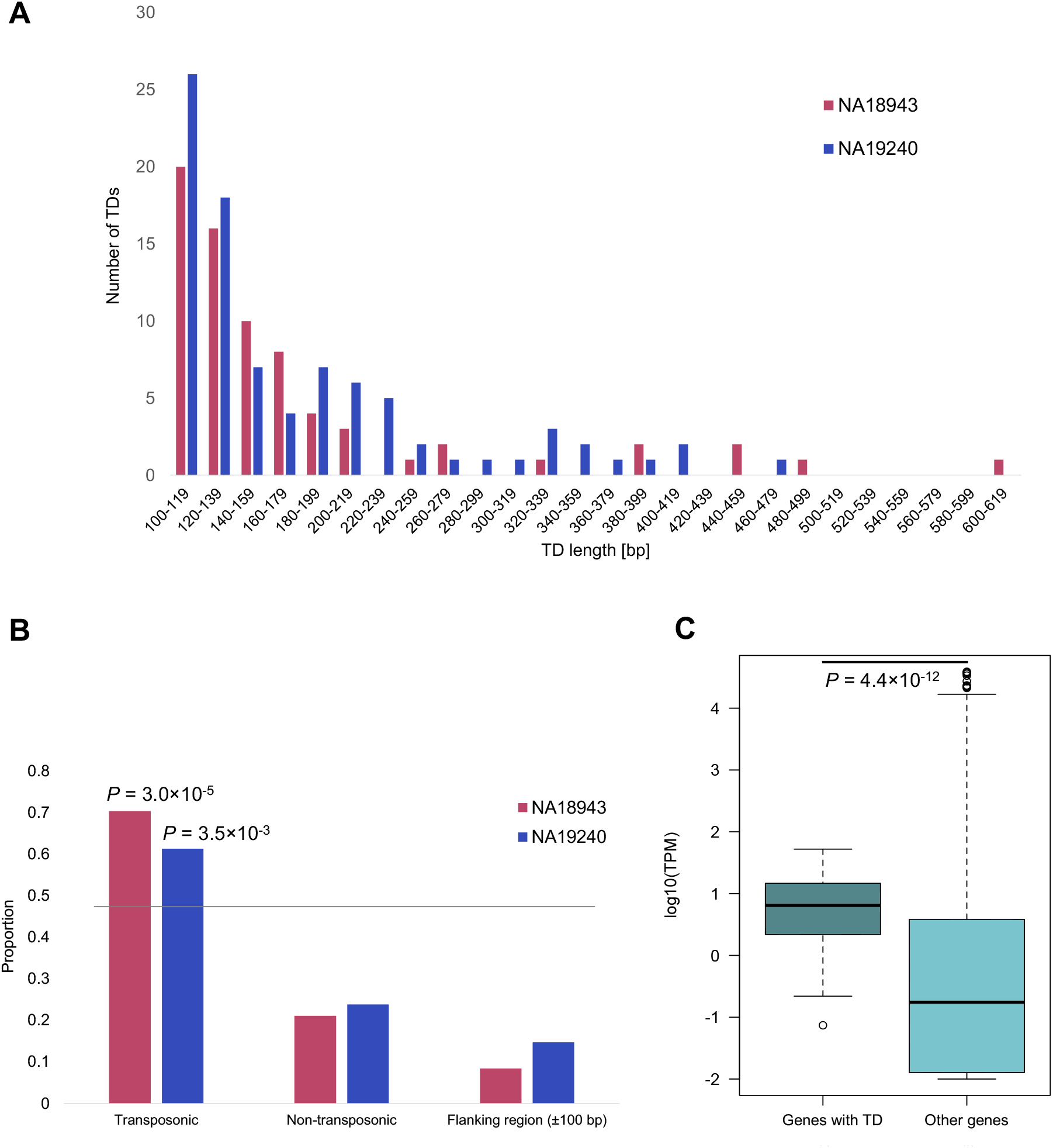
The characteristics of short TDs. **A**. The length distributions of TDs in NA18943 and NA19240. The numbers decreased as the length of TDs became longer. **B**. The enrichment of TDs in TEs. TDs are enriched in transposonic regions (79% in NA18943 and 76% in NA19240 when flanking regions are included). The grey line is the expected value of TDs in TE. **C**. The association of gene expression levels between TDs and genes with TD. Genes containing TD were expressed higher than other genes. TPM: Transcripts Per Million.

## Discussion

Assembling and polishing noisy long reads for target regions can be a useful approach for the investigation of SVs underlying the human genome. This study showed that LoMA can build accurate CSs from ONT data with substantial errors and revealed fine structures of insertions. We evaluated the accuracy of LoMA by a comparison with GRCh38 and found that LoMA reduced errors from 8.7% (raw ONT reads) to 0.76% (CS) (Fig. 1C). Sanger sequencing also estimated the error rate of CSs as 0.29% (Fig. 1C). Additionally, most heterozygous SVs detected by LoMA were successfully validated by PCR (Additional file 2: Fig. S1), suggesting a good accuracy of haplotype representation. We also compared the accuracy of CSs between LoMA and a tool with a similar function, lamassemble, finding LoMA has a lower average error rate (Fig. 1C). Notably, the error patterns showed different features: the insertion rate was larger in LoMA in many cases (69%, 74 from 108 regions), but the deletion rate was larger in lamassemble in most cases (94%, 102 from 108 regions) (Additional file 1: Table S1). These findings indicate that LoMA has the advantage of a lower deletion rate, although both tools may have their own systematic biases. Moreover, LoMA is superior in haplotype representation. Many *de novo* assemblers and lamassemble lack haplotype resolution, which results in representing one haplotype and a decline in accuracy [12][15]. Taken together, these results demonstrated that LoMA has sufficient accuracy to investigate genomic sequences at single-base resolution.

Since a large part of the human genome is identical within human populations [39], an analysis of unclear regions, which is typically manageable and quick, should be enough for most human genome studies. We focused on the regions full of clipped or collapsed reads (Fig. 2A), because such regions are likely to contain SVs and unstable structures. These unclear regions were scattered throughout the entire genome, but the clusters were observed in the pericentromeric and *HLA* region, as expected (Fig. 2B).

We applied LoMA to unclear regions. Generally, insertions are difficult to resolve because inserted sequences complicate alignments and remain uncharacterized without high-accuracy assembled sequences. We found 5,516 and 6,542 insertions (100 bp) in NA18943 and NA19240, respectively (Fig. 3B) and restored them from single-platform data. This analysis identified some interesting examples that showed the effectiveness of our approach. First, a processed pseudogene of *RPLP0* (Fig. 3D) was accurately aligned to the original gene on chromosome 12 (99.9% in identity), although *RPLP0* has many reverse-transcribed copies scattered on other chromosomes. Among them, the identified insertion was most accurately aligned to the original gene (specific splicing variant) rather than the other reverse-transcribed copies (the second most accurate alignment was 98.7% in identity), suggesting that the processed pseudogene was derived from the original *RPLP0* gene, not from other pseudogenes. This result indicates the importance of high-quality CSs for biological analyses. Second, we identified 15% of insertions of the alternative sequences in NA19240 were undetected in the standard SV set (Additional file 1: Table S6), which were caused by mapping errors in a previous study [24]. Particularly, we corrected the SV structure (Fig. 3E) in the *HLA* region, which is known for its hypervariability in human populations [40]. In NA18943, an insertion (∼14 kbp) derived from an alternative sequence was highly concordant (99.8% in identity) (Additional file 2: Fig. S2). Since such long insertions are difficult to restore by a mapping-based approach, this result indicates the efficiency of our assembly approach.

Our analysis showed that TR expansions were highly variable in length (Fig. 4A). Large TR expansions found in our analysis ranged to thousands of base pairs. Although the landscape of TR expansions has been studied using short and long reads [41][42][43], no studies have constructed long repeats and assessed the expansion rate throughout genomes. Many studies have identified pathogenic long repeat expansions [2][43], and several methods were developed to detect repeat expansions. However, such laboratory techniques are not suitable for multiple loci [43]. Thus, genome-wide analysis using LoMA will help explore TR expansions more widely and simply. Further, our analysis suggests that TR expansions exist even in healthy individuals, and these polymorphisms may affect disease susceptibility. Indeed, our result showed healthy individuals had expansions in exons consisting of TRs, and most of them were associated with various human traits and disease susceptibility (Additional file 1: Table S7) [44][45]. Further studies of human populations using long reads may identify disease-related TR expansions. We also observed TR expansions inside TEs, especially in SVAs (Fig. 4D), suggesting the existence of repetitive polymorphisms in repetitive sequences. A recent study reported SVs in SVAs associated with neurological disorders [46]. Our result is consistent with this previous study and indicates the importance of investigating TR expansions in TEs. This kind of variation (nested repeats) is hard to identify without long reads, and a localized assembly approach is necessary to interpret them. Therefore, investigations of human genetic variations using our method may lead to findings of novel susceptible genes of complex diseases. We also showed that TR expansions are prone to exist downstream of SINEs (*Alu* and *MIR*) (Fig. 4D). This relationship may reflect the reverse transcription mechanism in which an RNA element introduces a poly(A) sequence into the inserted site and provide a source of genomic instability around the site that gives rise to TR expansions.

*Alu* elements are a non-autonomous retrotransposons, and their insertion is initiated by target site-primed reverse transcription dependent on L1 endonuclease and reverse transcriptase, although there is no consensus on how the synthesized end is integrated to the target sequence on the opposite DNA strand [33][47]. We showed that the first nick induced by the initiation of the target-site primed reverse transcription of an *Alu* element has a strong motif, which is consistent with sites that L1 ORF2p endonuclease recognizes [48]; however, our analysis also suggested this recognition can take place at other sites that contain G-rich sequences (Fig. 5A). The length of a TSD is dependent on the second nicking site in the sense strand [33]. A past study suggested that TSD lengths are within 15-16 bp, though the investigated size of an *Alu* element was relatively small [49]. Our result corroborated that finding and further suggested that the distribution was skewed left, with an average length of 15 bp (Fig. 5B).

Lastly, we showed the characteristics of TDs. This class of insertions is understudied due to a lack of genome-wide sequence analysis at single-base resolution. We found that genes containing a TD were highly expressed compared to other genes (Fig. 6C), suggesting that TDs are induced by transcription stress. Furthermore, TDs were enriched in transposable elements (Fig. 6B). It is known that inversions are induced in L1 retrotransposition [50]. TE integration may also be related to the birth of novel TDs by some mechanism through the reverse transcription process. In the present study, we did not analyze indels shorter than 100 bp in length. However, the length distribution of TDs suggested that a number of TDs lie in short ranges (Fig. 6A) [28]. The numerical dominance of short TDs may contribute to genetic variations in the human population.

We revealed the true structures of insertions with high accuracy and inferred mechanisms for the insertions. However, our study has several limitations to be addressed in the future. First, LoMA classified haplotypes based on SVs, but SNVs and short indels were not taken into account. This is because current long reads have high sequencing error rates, and haplotype classification based on SNVs may cause computational errors. As sequencing technologies and basecalling accuracies improve, SNV-based haplotype reconstruction should be possible in the near future. Second, SV callings in low-coverage regions had low precision (Fig. 2C) due to high error rates in the current platforms. Improvements in sequencing error rates will enable reliable data from low-coverage regions. Third, in the genome-wide survey of unclear regions, we could not obtain satisfactory results in centromeric satellites and other complex regions, because the lengths of the reads were not sufficient to completely resolve these complex regions (Additional file 1: Table S2) [19]. Longer reads should make it possible to analyze these regions.

## Conclusions

Localized assembly using long reads is a promising approach to explore genetic variations in human populations. We demonstrated the effectiveness of our approach by revealing the true structures of insertions and repetitive regions at single-base resolution. The application of this approach to human disease studies will enable us to find novel pathogenic variants of Mendelian and complex disorders.

## List of abbreviations

ONT: Oxford Nanopore Technologies
PacBio: Pacific Biosciences
SV: Structural Variant
TR: Tandem Repeat
TE: Transposable Element
T2T: Telomere-to-Telomere
CS: Consensus Sequence
WGS: Whole-Genome Sequencing
TD: Tandem Duplication
NUMT: Nuclear Mitochondrial DNA sequence
TRF: Tandem Repeats Finder
UTR: Untranslated Region
STR: Short Tandem Repeat
TSD: Target Site Duplication
SD: Standard Deviation

## Declarations

### Ethics approval and consent to participate

Not applicable.

### Consent for publication

Not applicable.

### Availability of data and materials

The sequencing data have been deposited in the NBDC database in Japan under the accession number XXX (not yet uploaded) [51]. The source code can be downloaded from the author’s GitHub page: https://github.com/kolikem/loma [52].

### Authors’ contributions

KI developed LoMA and analyzed the sequencing data. HF and KI performed the experimental validations. KI and AF wrote the manuscript. All authors read and approved the final manuscript.

## Acknowledgements

We thank the Human Genome Center, the Institute of Medical Science, The University of Tokyo, for providing access to the SHIROKANE supercomputer. We also thank Dr. Wong Jing Hao, Dr. Yusuke Sano, Dr. Arinobu Fukunaga, and Ms. Kugui Yoshida for great assistance.

## URLs

European Nucleotide Archive. https://www.ebi.ac.uk/ena/browser/home. Accessed 27 September 2022.

Guppy. https://community.nanoporetech.com/downloads. Accessed 27 September 2022.

A.F.A. Smit, R. Hubley & P. Green RepeatMasker at http://repeatmasker.org. Accessed 27 September 2022.

RefSeq. https://www.ncbi.nlm.nih.gov/refseq/rsg/. Accessed 27 September 2022. GTEx data. https://gtexportal.org/home/. Accessed 27 September 2022.

UCSC Genome Browser. https://genome.ucsc.edu. Accessed 27 September 2022.

URL6. The NBDC database. https://humandbs.biosciencedbc.jp. Accessed 28 September 2022. URL7. Ko Ikemoto’s GitHub. https://github.com/kolikem/. Accessed 28 September 2022.

## Supplementary Information

**Additional file 1:** Supplementary Tables.

**Additional file 2:** Supplementary Figures and information.

## References

1. Logsdon GA, Vollger MR, EichlerE E. Long-read human genome sequencing and its applications. Nat Rev Genet. 2020;21:597–614. http://dx.doi.org/10.1038/s41576-020-0236-x.

2. Sone J, Mitsuhashi S, Fujita A, Mizuguchi T, Hamanaka K, Mori K, et al. Long-read sequencing identifies GGC repeat expansions in NOTCH2NLC associated with neuronal intranuclear inclusion disease. Nat Genet. 2019;51:1215–21. http://dx.doi.org/10.1038/s41588-019-0459-y.

3. Fujimoto A, Wong JH, Yoshii Y, Akiyama S, Tanaka A, Yagi H, et al. Whole-genome sequencing with long reads reveals complex structure and origin of structural variation in human genetic variations and somatic mutations in cancer. Genome Med. 2021;13:1–15. https://doi.org/10.1186/s13073-021-00883-1.

4. Miga KH, Koren S, Rhie A, Vollger MR, Gershman A, Bzikadze A, et al. Telomere-to-telomere assembly of a complete human X chromosome. Nature. 2020;585:79–84. http://dx.doi.org/10.1038/s41586-020-2547-7.

5. Sano Y, Koyanagi Y, Wong JH, Murakami Y, Fujiwara K, Endo M, et al. Likely pathogenic structural variants in genetically unsolved patients with retinitis pigmentosa revealed by long-read sequencing. J Med Genet. 2022;jmedgenet-2022-108428. https://doi.org/10.1136/jmedgenet-2022-108428.

6. treangen TJ, SalzbergS L. Repetitive DNA and next-generation sequencing: Computational challenges and solutions. Nat Rev Genet. 2012;13:36–46. https://doi.org/10.1038/nrg3117.

7. tang H, Kirkness EF, Lippert C, Biggs WH, Fabani M, Guzman E, et al. Profiling of Short-Tandem-Repeat Disease Alleles in 12,632 Human Whole Genomes. Am J Hum Genet. 2017;101:700–15. https://doi.org/10.1016/j.ajhg.2017.09.013.

8. HannanA J. Tandem repeats mediating genetic plasticity in health and disease. Nat Rev Genet. 2018;19:286–98. http://dx.doi.org/10.1038/nrg.2017.115.

9. Payer LM, Steranka JP, Yang WR, Kryatova M, Medabalimi S, Ardeljan D, et al. Structural variants caused by Alu insertions are associated with risks for many human diseases. Proc Natl Acad Sci U S A. 2017;114:E3984–92. https://doi.org/10.1073/pnas.1704117114.

10. Mavragani CP, Sagalovskiy I, Guo Q, Nezos A, Kapsogeorgou EK, Lu P, et al. Expression of Long Interspersed Nuclear Element 1 Retroelements and Induction of Type I Interferon in Patients With Systemic Autoimmune Disease. Arthritis Rheumatol. 2016;68:2686–96. https://doi.org/10.1073/pnas.1704117114.

11. Sergey Nurk, Sergey Koren AR et al. The complete sequence of a human genome. Science. 2022;376:44–53. https://doi.org/10.1126/science.abj6987.

12. Mahmoud M, Gobet N, Cruz-Dávalos DI, Mounier N, Dessimoz C, Sedlazeck FJ. Structural variant calling: The long and the short of it. Genome Biol. 2019;20:1–14. https://doi.org/10.1186/s13059-019-1828-7.

13. Nattestad M, SchatzM C. Assemblytics: A web analytics tool for the detection of variants from an assembly. Bioinformatics. 2016;32:3021–3. https://doi.org/10.1093/bioinformatics/btw369.

14. Baker M. De novo genome assembly: what every biologist should know. Nature Methods. 2012;9:333–337. https://doi.org/10.1038/nmeth.1935.

15. Frith MC, Mitsuhashi S, Katoh K. lamassemble: Multiple Alignment and Consensus Sequence of Long Reads. In: Katoh K, editor. Mult Seq Alignment Methods Protoc. 2021;2231:135–145. https://doi.org/10.1007/978-1-0716-1036-7_9.

16. Li H. Minimap2: Pairwise alignment for nucleotide sequences. Bioinformatics. 2018;34:3094–100. https://doi.org/10.1093/bioinformatics/bty191.

17. Katoh K, StandleyD M. MAFFT multiple sequence alignment software version 7: Improvements in performance and usability. Mol Biol Evol. 2013;30:772–80. https://doi.org/10.1093/molbev/mst010.

18. Belmont JW, Hardenbol P, Willis TD, Yu F, Yang H, Ch’Ang LY, et al. The International HapMap Project. Nature. 2003;426:789–96. https://pubmed.ncbi.nlm.nih.gov/14685227.

19. De Coster W, De Rijk P, De Roeck A, De Pooter T, D’Hert S, Strazisar M, et al. Structural variants identified by Oxford Nanopore PromethION sequencing of the human genome. Genome Res. 2019;29:1178–87. https://doi.org/10.1101/gr.244939.118.

20. European Nucleotide Archive. https://www.ebi.ac.uk/ena/browser/home. Accessed 27 September 2022.

21. Fujimoto A, Nakagawa H, Hosono N, Nakano K, Abe T, Boroevich KA, et al. Whole-genome sequencing and comprehensive variant analysis of a Japanese individual using massively parallel sequencing. Nat Genet. 2010;42:931–6. https://doi.org/10.1038/ng.691.

22. Guppy. https://community.nanoporetech.com/downloads. Accessed 27 September 2022.

23. Li H, Handsaker B, Wysoker A, Fennell T, Ruan J, Homer N, et al. The Sequence Alignment/Map format and SAMtools. Bioinformatics. 2009;25:2078–9. https://doi.org/10.1093/BIOINFORMATICS/BTP352.

24. ChaissonM J. Multi-platform discovery of haplotype-resolved structural variation in human genomes. Nat Commun. 2019;10:1784. https://doi.org/10.1038/s41467-018-08148-z.

25. Delage WJ, Thevenon J, Lemaitre C. Towards a better understanding of the low recall of insertion variants with short-read based variant callers. BMC Genomics. 2020;21:1–17. https://doi.org/10.1186/s12864-020-07125-5.

26. Benson G. Tandem repeats finder: a program to analyze DNA sequences. Nucleic Acids Res. 1999;27:573–80. https://academic.oup.com/nar/article/27/2/573/1061099.

27. A.F.A. Smit, R. Hubley & P. Green RepeatMasker at http://repeatmasker.org. Accessed 27 September 2022.

28. Ashouri S, Wong JH, Nakagawa H, Shimada M, Tokunaga K, Fujimoto A. Characterization of intermediate-sized insertions using whole-genome sequencing data and analysis of their functional impact on gene expression. Hum Genet. 2021;140:1201–16. https://doi.org/10.1007/s00439-021-02291-2.

29. KentW J. BLAT---The BLAST-Like Alignment Tool. Genome Res. 2002;12:656–64. https://doi.org/10.1101/gr.229202.

30. Ellegren H. Heterogeneous mutation processes in human microsatellite DNA sequences. Nat Genet. 2000;24:400–2. https://doi.org/10.1038/74249.

31. RefSeq. https://www.ncbi.nlm.nih.gov/refseq/rsg/. Accessed 27 September 2022.

32. BurnsK H. Transposable elements in cancer. Nat Rev Cancer. 2017;17:415–24. http://dx.doi.org/10.1038/nrc.2017.35

33. Deininger P. Alu elements: Know the SINEs. Genome Biol. 2011;12:1–12. https://doi.org/10.1186/gb-2011-12-12-236.

34. Bailey TL, Johnson J, Grant CE, Noble WS. The MEME Suite. Nucleic Acids Res. 2015;43:W39–49. https://doi.org/10.1093/nar/gkv416.

35. GTEx data. https://gtexportal.org/home/. Accessed 27 September 2022.

36. Altshuler DM, Durbin RM, Abecasis GR, Bentley DR, Chakravarti A, Clark AG, et al. An integrated map of genetic variation from 1,092 human genomes. Nature. 2012;491:56–65. https://doi.org/10.1038/nature11632.

37. Batzer MA, Deininger PL. Alu repeats and human genomic diversity. Nat Rev Genet. 2002;3:370–9. https://doi.org/10.1038/nrg798.

38. UCSC Genome Browser. https://genome.ucsc.edu. Accessed 27 September 2022.

39. Pang AW, MacDonald JR, Pinto D, Wei J, Rafiq MA, Conrad DF, et al. Towards a comprehensive structural variation map of an individual human genome. 2010;11. https://doi.org/10.1186/gb-2010-11-5-r52.

40. Shiina T, Hosomichi K, Inoko H, Kulski JK. The HLA genomic loci map: Expression, interaction, diversity and disease. J Hum Genet. 2009;54:15–39. https://doi.org/10.1038/jhg.2008.5.

41. Mitsuhashi S, Frith MC, Mizuguchi T, Miyatake S, Toyota T, Adachi H, et al. Tandem-genotypes: robust detection of tandem repeat expansions from long DNA reads. Genome Biol. 2019;20:1–17. https://doi.org/10.1186/s13059-019-1667-6.

42. trost B, Engchuan W, Nguyen CM, Thiruvahindrapuram B, Dolzhenko E, Backstrom I, et al. Genome-wide detection of tandem DNA repeats that are expanded in autism. Nature. 2020;586:80–6. http://dx.doi.org/10.1038/s41586-020-2579-z.

43. Chintalaphani SR, Pineda SS, Deveson IW, Kumar KR. An update on the neurological short tandem repeat expansion disorders and the emergence of long-read sequencing diagnostics. Acta Neuropathol Commun. 2021;9. https://doi.org/10.1186/s40478-021-01201-x.

44. Kim HS, Lyons KM, Saitoh E, Azen EA, Smithies O, Maeda N. The structure and evolution of the human salivary proline-rich protein gene family. Mamm Genome. 1993;4:3–14. https://doi.org/10.1007/BF00364656.

45. Mangino M, Hwang SJ, Spector TD, Hunt SC, Kimura M, Fitzpatrick AL, et al. Genome-wide meta-analysis points to CTC1 and ZNF676 as genes regulating telomere homeostasis in humans. Hum Mol Genet. 2012;21:5385–94. https://doi.org/10.1093/hmg/dds382.

46. van Bree EJ, Guimarães RLFP, Lundberg M, Blujdea ER, Rosenkrantz JL, White FTG, et al. A hidden layer of structural variation in transposable elements reveals potential genetic modifiers in human disease-risk loci. Genome Res. 2022;32:656–70. https://doi.org/10.1101/gr.275515.121.

47. Chen JM, Férec C, Cooper DN. Mechanism of Alu integration into the human genome. Genomic Med. 2007;1:9–17. https://doi.org/10.1007/s11568-007-9002-9.

48. Feng Q, Moran J V., Kazazian HH, Boeke JD. Human L1 retrotransposon encodes a conserved endonuclease required for retrotransposition. Cell. 1996;87:905–16. https://doi.org/10.1016/S0092-8674(00)81997-2.

49. Jurka J. Sequence patterns indicate an enzymatic involvement in integration of mammalian retroposons. Proc Natl Acad Sci U S A. 1997;94:1872–7. https://doi.org/10.1073/pnas.94.5.1872.

50. Ostertag EM, Kazazian J. Twin priming: A proposed mechanism for the creation of inversions in L1 retrotransposition. Genome Res. 2001;11:2059–65. https://doi.org/10.1101/gr.205701.

51. The NBDC database. https://humandbs.biosciencedbc.jp. Accessed 28 September 2022.

52. Ko Ikemoto’s GitHub. https://github.com/kolikem/. Accessed 28 September 2022.

