## Additional_file_2 for "Localized assembly for long reads enables genome-wide analysis of repetitive regions at single-base resolution in human genomes"

### Supplementary Materials

#### Counting STR types

In principle, it is difficult to distinguish in which strand a TR expansion event happened by observing DNA sequences. Additionally, it is also difficult to analyze STRs tandemly repeated many times after discriminating sorted sequences that keep their order (e.g., CGA, ACG and GAC). Therefore, we treated and counted a group of STRs that were the same in lexicographical order without discriminating strands as the same group.

#### Genome-wide analysis of SVs using LoMA

The success rate of the reconstruction of unclear regions was 88.1% and 92.2% in NA18943 and NA19240, respectively, excluding the failed regions derived from centromeric regions. In NA18943, 3,065 regions were classified as heterozygous, and 10,757 were homozygous regions. In NA19240, 7,542 regions were heterozygous, and 7,678 were homozygous. The difference in ratio of heterozygous and homozygous regions reflected the high genetic diversity in the African population. We then assessed the success rate of LoMA in regions in which clipped sequences accumulated, because sequence information is not accessible in regions full of clipped reads, and the clipped sequences are ignored from typical analyses. In NA18943 and NA19240, 82.5% and 71.8%, respectively, of the clipped-sequence-accumulating regions were correctly assembled excluding centromeres.

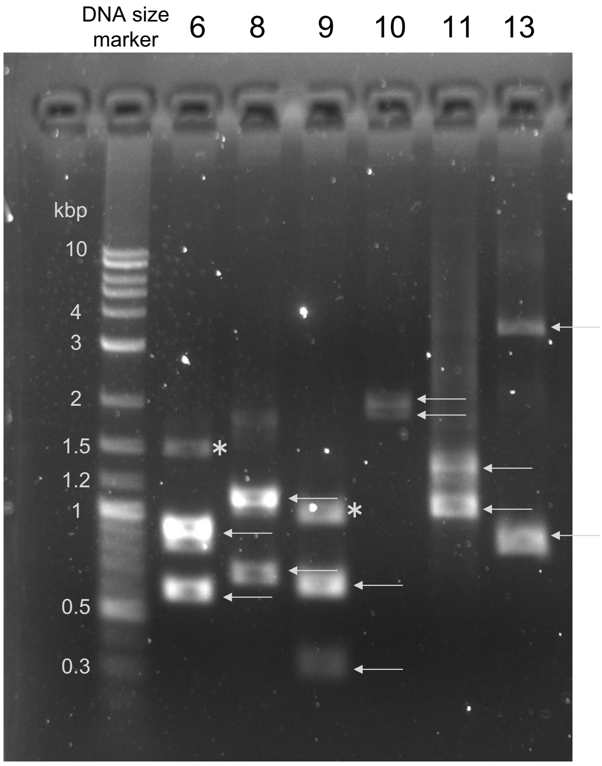


**Fig. S1:** Examples of PCR validation. Six from eight heterozygous variants validated by PCR amplification are shown in the picture of electropherogram. The numbers are the variant IDs in Table S4. White arrows indicate the amplified products in expected sizes. The asterisks are non-specific bands.

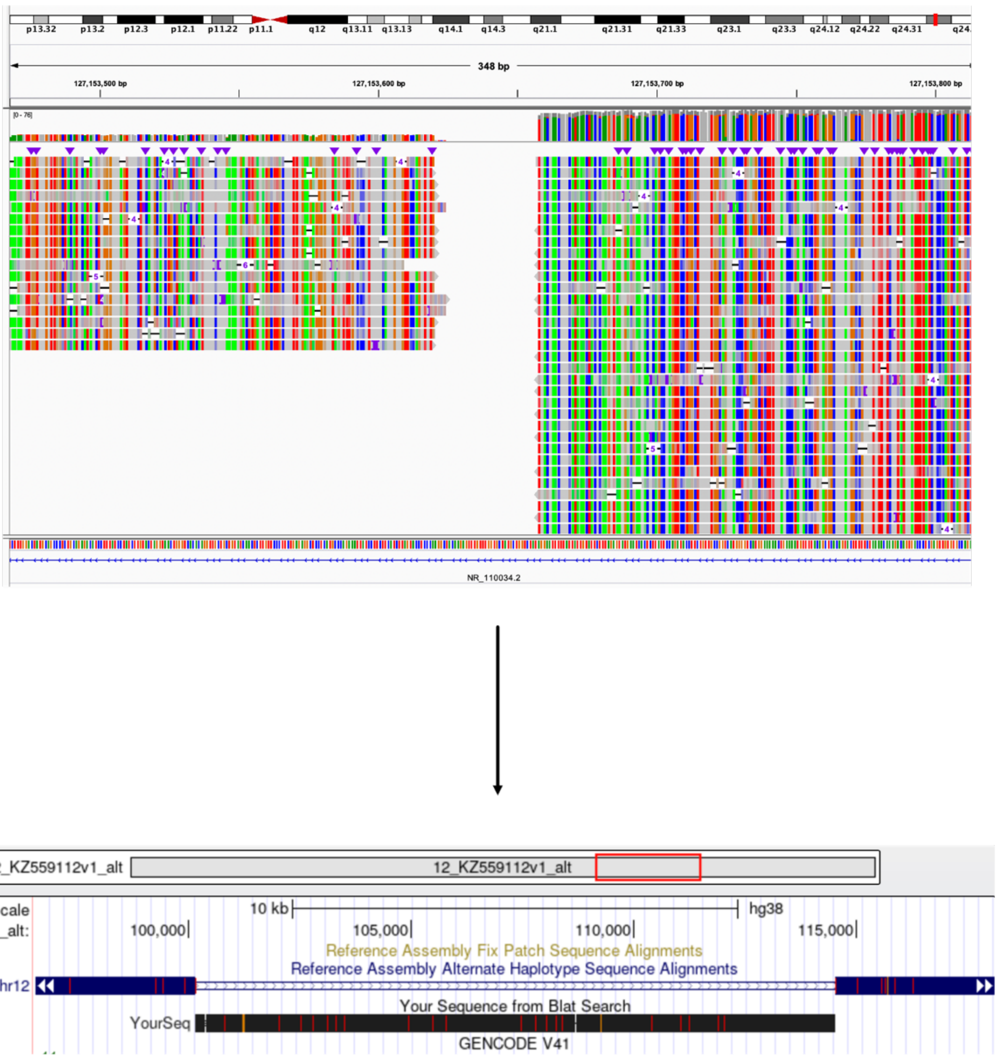


**Fig. S2:** The longest alternative sequence insertion. A long insertion (~14 kbp) that accurately mapped (99.8% in identity) to an alternative sequence on chr12 was detected in NA18943. It showed the clip-accumulated pattern as visualized by IGV [1]. After the restoration of this region, the sequence was clearly mapped to the alternative sequence on Genome Browser [2].

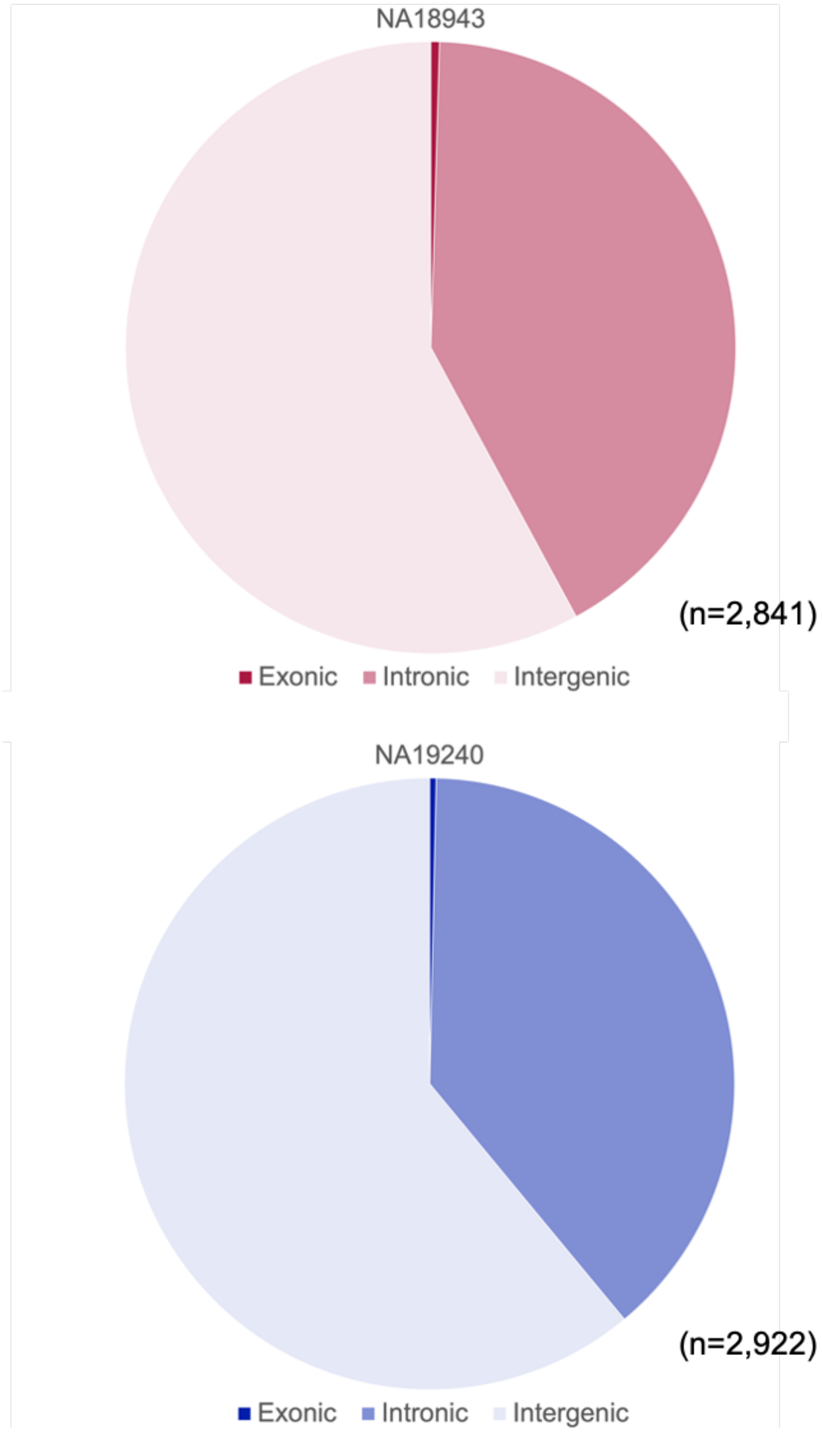


**Fig. S3** Genic and intergenic TR expansions. In NA18943, genic TDs accounted for 42%, and in NA19240 they were 39%, which is consistent with the size of the genic region in the human genome. Only 7 and 12 genes contained exonic TR expansions in NA18943 and NA19240, respectively.

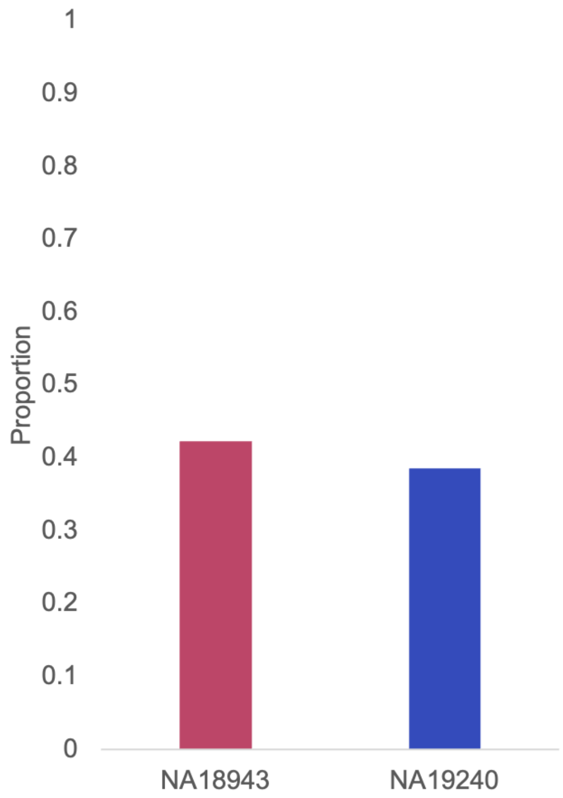


**Fig. S4** Genic TDs. In NA18943, 30 TDs (42%) occurred in the genic region (all in non-CDS regions); in NA19240, it was 34 TDs (39%; all in non-CDS regions). These observations are consistent with the size of the genic region in the human genome. TDs were not enriched in genes.

**References**

1. Thorvaldsdóttir H, Robinson JT, Mesirov JP. Integrative Genomics Viewer (IGV): High-performance genomics data visualization and exploration. Brief Bioinform. 2013;14:178–92. https://doi.org/10.1093/bib/bbs017.

2. Genome Browser. https://genome.ucsc.edu/index.html. Accessed 17 October 2022.
